# F-Seq2: improving the feature density based peak caller with dynamic statistics

**DOI:** 10.1101/2020.10.06.328674

**Authors:** Nanxiang Zhao, Alan P. Boyle

**Affiliations:** Department of Computational Medicine and Bioinformatics, University of Michigan, Ann Arbor, MI 48109, USA; Department of Human Genetics, University of Michigan, Ann Arbor, MI 48109, USA

## Abstract

Genomic and epigenomic features are captured at a genome-wide level by using high-throughput sequencing technologies. Peak calling is one of the first essential steps in analyzing these features by delineating regions such as open chromatin regions and transcription factor binding sites. Our original peak calling software, F-Seq, has been widely used and shown to be the most sensitive and accurate peak caller for DNase I hypersensitive sites sequencing (DNase-seq) data. However, F-Seq lacks support for user-input control dataset nor reporting test statistics, limiting its ability to capture systematic and experimental biases and accurately estimate background distributions. Here we present an improved version, F-Seq2, which combined the power of kernel density estimation and a dynamic “continuous” Poisson distribution to robustly account for local biases and solve ties when ranking candidate peaks. In F-score and motif distance analysis, we demonstrated the superior performance of F-Seq2 than other competing peak callers used by the ENCODE Consortium on simulated and real ATAC-seq and ChIP-seq datasets. The output of F-Seq2 is suitable for irreproducible discovery rate (IDR) analysis as the test statistics calculated for individual candidate summit and ties are robustly solved.

## INTRODUCTION

High-throughput sequencing (HTS) has become a central technology in deciphering genomic and epigenomic landscapes. Various assays exist to identify specific biological events. DNase I hypersensitive sites sequencing (DNase-seq) (1), Assay of Transposase Accessible Chromatin sequencing (ATAC-seq) (2), and Formaldehyde-Assisted Isolation of Regulatory Elements sequencing (FAIRE-seq) (3) are designed to detect genome-wide chromatin accessibility. Transcription factor (TF) binding and histone modifications are measured using Chromatin Immuno-Precipitation sequencing (ChIP-seq) (4, 5). The short read sequences produced by these assays are usually filtered and mapped back to a reference genome, then accumulated and piled up in genomic regions. The enrichment (e.g. counts) of mapped reads can be abstractly viewed as a digital signal of relevant biological events varying along the genome. The genome-wide enrichment signal can be further processed with a peak-calling program, or peak caller, to find the arguments of local maxima (argmax), representing discrete loci with statistically significant enrichment over background for the relevant biological event. For example, individual TF binding sites in a ChIP-seq experiment.

We introduced F-Seq as a general peak caller for DNase-seq and ChIP-seq in 2008 (6). Unlike other recent methods (7, 8), F-seq calls peaks in HTS signals by reconstructing the genome-wide signal using a kernel density estimator (KDE) (9, 10). KDE-based reconstructed signal is smoother and more accurate than histogram-based methods (e.g. sliding window), but still interpretable and useful for visualization as the estimate is proportional to the probability of finding a read at a given base pair (11). A Gaussian kernel with a chosen bandwidth is centered at each read and kernels are summed up to obtain the density estimate. Peak regions are then called if the signal is higher than the threshold calculated from a simulated background model. F-Seq has been widely used in the ENCODE project (12) and beyond, which is shown to be more accurate and sensitive than competing peak callers for DNase-seq data (13). However, F-Seq lacks native support for a separate control dataset. Consequently, F-Seq cannot capture or eliminate local biases affecting read distribution along the genome, such as copy number variation, read mappability, and local chromatin structure (7). This limits the performance of F-Seq especially on ChIP-seq data since the majority of ChIP-seq experiments have corresponding control data which contains unique information for accurate peak calling (14). In addition, F-Seq does not report tests statistics (e.g. p-value or q-value) apart from the signal value at each position.

To address these shortcomings, we have developed F-Seq version 2 (F-Seq2), a complete rewrite of the original F-Seq in Python. F-Seq2 implements a dynamic parameter to conduct local statistical analysis with an underlying “continuous” Poisson distribution. By combining the power of the local test and the KDE, which model the read probability distribution with statistical rigor, we robustly account for local biases and solve ties that occur when ranking candidate summits, making results suitable for irreproducible discovery rate (IDR) analysis (15). We compared F-Seq2 with four peak callers used by the ENCODE Consortium (12) on simulated and real ChIP-seq and ATAC-seq datasets, demonstrating performance gains arising from the joint effect of KDE and the local test, especially in the absence of control data.

## MATERIAL AND METHODS

### Density profiles and peak calling

The original F-Seq KDE was modified to use the unnormalized density estimates, where the sum is limited to the number of reads proximal to any given position for computational convenience for statistical value calculation. Scaling between control and treatment data was necessary when sequencing depths were different. Control data was scaled to treatment data at an individual chromosome level as the ratios of their total reads fluctuated among different chromosomes. The reconstructed signal by KDE was treated as a digital signal emitted on a chromosome. Argmax of the signal (i.e. the candidate summits for statistical testing) were determined using the following criteria: minimum height, distance and prominence. Estimation of the fragment size for ChIP-seq data and the simulated background threshold for defining and selecting candidate summits and delineating final peak regions were unchanged from the original F-Seq method.

We adopted and modified the dynamic testing idea introduced by MACS2 (7) to assign each candidate summit a statistical enrichment value related to a background. Rather than using a constant background estimation for all candidates, a local background was estimated for each candidate providing a more accurate method to calculate enrichment p-values due to the local fluctuations of read enrichment distributions. The Poisson distribution (characterized by Λ) was used to model the number of reads from a genomic region as this is mathematically proven more powerful compared to the Binomial distribution in peak calling (16). Specifically, *λ* for a summit is defined as *λ_local_ = max*(*λ_BG_*, [*λ*_*p*1_, *λ*_1*k*_], *λ*_5*k*_, *λ*_10*k*_), where *λ*_*p*1_ is the maximum signal value for one pseudo-read, *λ_BG_* is the estimate of the individual chromosome background, and *λ_x_* is the estimate of a *x* bp window centered at the summit. All the estimations are from control data where available. Otherwise, estimates were taken from treatment data and small regions (square brackets in the formula) were excluded to alleviate the background estimation boost by the summit signal value. Since Poisson distribution is a discrete probability distribution and signals are continuous values, the calculation of probability mass function (*pmf*) of a float that lies between two integers requires interpolation. The *pmf* was transformed to the logarithmic space, where the *pmf* of a float is linearly interpolated between two adjacent integers. The interpolation in log space solves ties in peak ranks by transforming the Poisson into the “continuous” distribution. Multi-test correction was then conducted with the Benjamini-Hochberg approach (17) to calculate q-values (more precisely, false discovery rate adjusted p-values) from the interpolated p-values.

### Benchmarking with selected peak callers

Four peak callers were selected along with F-Seq2 to benchmark on 100 simulated HTS datasets, 3 real ChIP-seq datasets, and one ATAC-seq dataset. The comparison methods which are used by the ENCODE included Model-based Analysis for ChIP-Seq version 2 (MACS2) (7), SPP (18), MUltiScale enrIchment Calling for ChIP-Seq (MUSIC) (8), and Genome wide Event finding and Motif discovery (GEM) (19). 100 treatment and paired control samples were simulated to closely resembles real ChIP-seq data which were reproduced from (16), allowing us to evaluate peak callers under different scenarios in a situation where the ground truth is known. Real ChIP-seq datasets were obtained from ENCODE where 3 different TFs are in different cell lines (12). As the ground truth is unknown in real datasets, one common alternative is to use TF binding motifs. Motifs were obtained from the JASPAR database (20) irrespective of cell line specificity and used for the 3 real ChIP-seq datasets. Similarly, the union of 117 TF ChIP-seq conservative IDR peaks from ENCODE were used as the “ground truth” for the ATAC-seq benchmarking. Raw ATAC-seq bam files were downloaded from (2) (see Availability for data accession numbers).

Performance for all peak callers was evaluated across a range of significance thresholds representing the different number of top ranked peaks. The main evaluation metric was F-score, wherein a higher F-score indicates a more balanced performance in terms of precision and recall. All peak callers were run with recommend settings and the lowest possible (most conservative) thresholds (see Availability for parameters settings).

### Evaluation for simulated data

Typically, peak calling results are not directly comparable as they have different peak widths and different estimated p-values or q-values generated from different statistical tests. To address this issue, all tools were first run with the lowest possible threshold to obtain an extensive list of peaks on each simulated dataset for each tool. Then all peaks were limited to a 200 bp window centered at the peak summit or peak centers depending on the information available. Operating characteristics can be evaluated by varying the threshold to obtain the same top number of peaks from each tool, where peaks are ranked by individual significance measurements. F-score was used as the evaluation metric, which is the harmonic mean of precision and recall. True positives were defined as the number of predicted peaks which overlap with ground truth peaks, precision as the fraction of true positive peaks in all predictions, and recall as the fraction of true positive peaks in all ground truth peaks. A higher F-score indicates a more balanced and optimal performance in terms of both precision and recall.

### Evaluation for ATAC-seq data

Evaluation of F-Seq2 and MACS2 used the union of 117 conservative IDR peaks of TF ChIP-seq from ENCODE as the “ground truth”. All IDR peaks were in the GM12878 cell line to be comparable to the ATAC-seq data. Each tool was run with two main modes, single-end (SE) and paired-end (PE) mode. Paired-end mode has the advantage of knowing the exact fragment length, which is useful when filtering out fragments whose length fall within a certain range to avoid peak calls on nucleosome centers (2). Operating characteristics curves were plotted similarly as in evaluation for simulation data by the varying respective thresholds. The main difference was that the metric was changed to F0.5-score and with new definitions. We used F0.5-score to put more emphasis on precision than recall due to the incompleteness of our “ground truth”. True positives were redefined as the number of base pairs (bp) of the predicted peaks that overlap with ground truth peaks, precision as the fraction of the correctly predicted base pairs in all predictions, and recall as the fraction of the correctly predicted base pairs in all ground truth peaks. The reason for the new definitions is that ATAC-seq peak lengths are usually larger than TF ChIP-seq peak lengths. We shifted focus from evaluating on summits to peak regions for a more comprehensive evaluation.

### Evaluation for real TF ChIP-seq data

Evaluations of real TF ChIP-seq peak calling results required additional datasets, JASPAR motif Position Weight Matrices (PWM) of each TF. Motif positions were identified by TFM P-value and Bowtie programs (21, 22) scanning along the genome with the PWMs. For each ChIP-seq dataset, each tool called a list of significant peaks with their default thresholds. The shortest distances between the significant peaks and the motifs of the corresponding TF were obtained and used as the main evaluation metrics. Specifically, we evaluated the fraction of top *n* up to 1000 peaks ranked by significance within a 100 bp window of a motif. We also examined the empirical cumulative distribution of the shortest distance of those top 1000 peaks for each tool.

### F-Seq2 auto filter design for paired-end ATAC-seq data peak calling

We designed the PE filter based on information in (2), where fragment length under 100 bp, between 180 and 247 bp, between 315 and 473 bp, and between 558 and 615 bp were considered nucleosome free, mono-, di-, and tri-nucleosomes, respectively. The difference between our analysis and Buenrostro’s (2), in which they only used fragments under 100 bp for open chromatin analysis, is that we included more fragments. By not including the ranges between the non-overlapping cutoffs, a large percentage (~15%) of fragments are discarded, leading to a reduction in recall. These fragments (e.g. between 100 and 180 bp) may contain useful information for identifying open chromatin regions (23). F-Seq2 takes advantage of more available reads to accurately estimate background distribution. Specifically, we only excluded fragments within mono-, di-, and tri-nucleosomes ranges. Fragments larger than 558 bp (i.e. multinucleosome-sized fragments) are also excluded as these fragments are associated with condensed heterochromatin (2).

## RESULTS

### Performance on simulated datasets

To accurately evaluate the peak callers under a variety of scenarios, each method was benchmarked on 100 pairs of simulated data. F-Seq2 and MACS2 were found to be the top two performers shown by the F-score operating characteristic curve (Figure 1A). The highest F-scores estimated by generalized additive models across 100 pairs were 0.897, and 0.884 for F-Seq2 and MACS2, respectively. Both methods outperformed MUSIC, the third-best method, by a margin of ~0.1 (MUSIC 0.781). Despite differences in implementing a dynamic parameter between F-Seq2 and MACS2, the performance gap suggests using a dynamic parameter in ranking peaks, effectively removing false positives, is a huge advantage, consistent with the conclusion from (16). The number of peaks called by the default threshold of each peak caller was compared to the number of peaks in the ground truth (Figure 1B). F-Seq2 best correlated with the ground truth (*r*=0.88). MUSIC (*r*=0.74) was slightly better than MACS2 (*r*=0.70). This high correlation indicates default threshold of F-Seq2 is reliable when estimating the number of significant peaks under a simulation setting.

**Figure 1.**
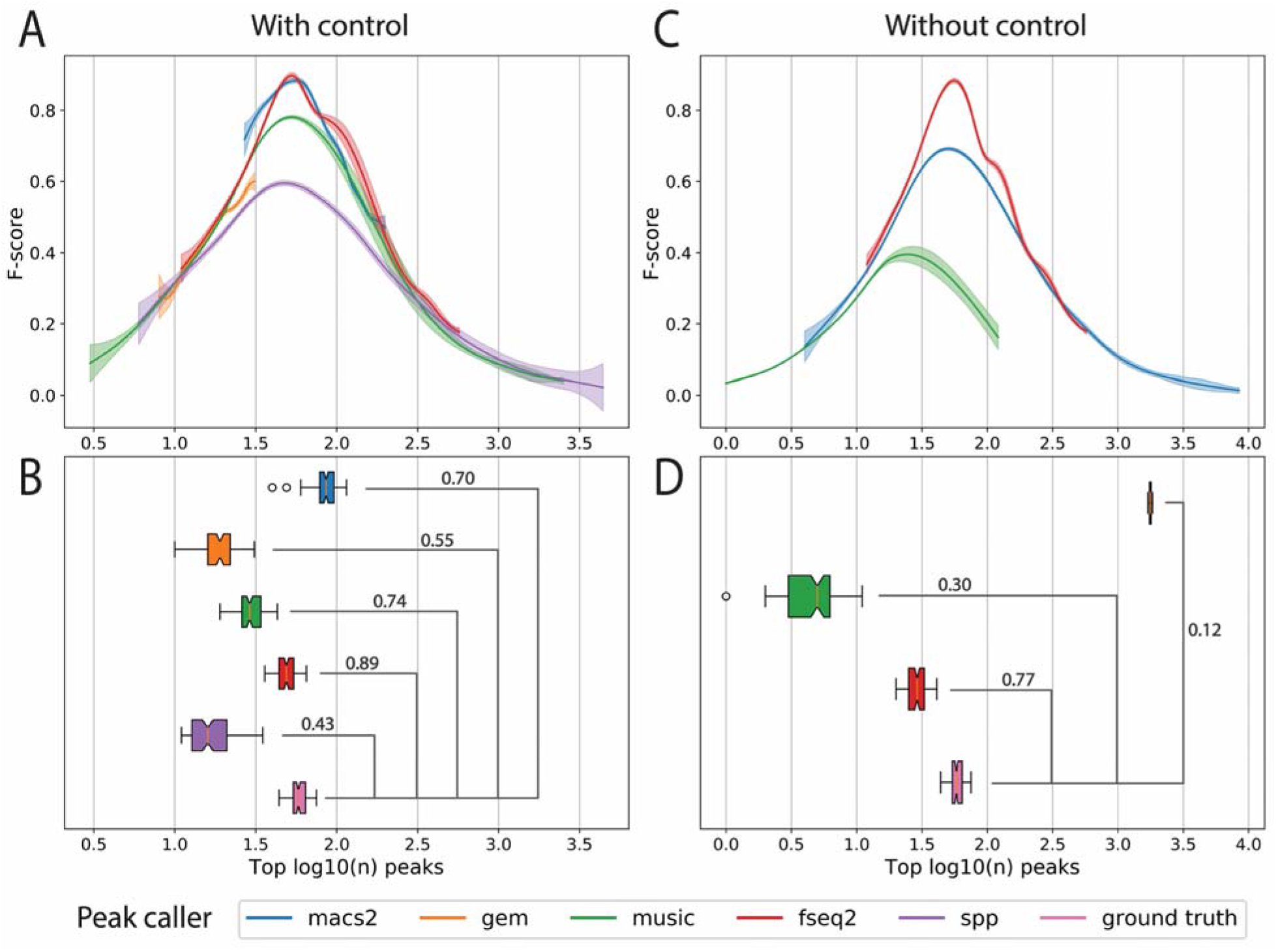
Comparison of peak callers on 100 pairs of simulated transcription factor ChIP-seq datasets. (A) The F-score operating characteristic curve where F-score is plotted as a function of the log_10_ top number of peaks called with control data. Generalized additive models are used to estimate the mean and 95% confidence intervals (shaded areas) of 100 peak calling results for each peak caller. (B) Boxplot of the number of peaks called by each peak caller with default threshold with control data, and the number of significant peaks in ground truth. Numbers are shown in log_10_ scale. Pearson’s correlation coefficient *r* is shown above the bridge linking peak caller and ground truth. (C) The F-score plot without control data. SPP was not able to run without control. GEM resulted in few peaks which is not shown in the plot. (D) Boxplot without control data.

Although control data is often essential for modeling background distributions for candidate summits, F-Seq2 demonstrated highly balanced performance between precision and recall on simulated ChIP-seq data without controls (Figure 1C&D). F-Seq2 stood out among the other peak callers and achieved comparable performance (0.883) to those with control datasets (0.897). These results suggest that the treatment data alone may contain a significant amount of control information at a large scale. This is also evident in real ChIP-seq datasets for FoxA1 ChIP-seq (7), where control read counts were correlated well with treatment read counts in 10 kb windows across the genome. This observed high correlation and the performance of F-seq2 implies that control information can be robustly extracted from treatment data and can used to estimate background distribution for peak calling, given it does not greatly contradict with the treatment data and given a statistically rigorous modeling method for treatment data (e.g. F-Seq2 KDE). For real ChIP-seq datasets, especially where the correlation is low between control and treatment data, calling peaks without control data is less accurate due to the loss of unique information within this dataset and cannot be recovered from treatment data (14).

### Performance on real datasets

The absence of control data is more often seen in DNase-seq and ATAC-seq experiments compared to ChIP-seq. Therefore, F-seq2 was directly compared to MACS2 on ATAC-seq data to further evaluate performance without control data (Figure 2). Both F-Seq2 with paired-end (PE) auto mode and MACS2 with single-end (SE) shift-extend mode, two different strategies to avoid calling peaks on nucleosome centers, precisely identified open chromatin regions with their top ranked peaks (see Material and Methods for auto filter design details). The better characteristic curve of F-Seq2 (highest F-0.5 score = 0.62) indicates the filter-based method is more effective in avoiding peaks called on nucleosomes compared to the shift-based method. MACS2 SE shift-extend mode outperformed its PE mode (highest F-0.5 scores: 0.58 vs. 0.54) at low genome coverage (1% of human genome). This precision gained by the shift-extend strategy is likely why single-end data is used as part of the official ENCODE ATAC-seq data analysis pipeline (12). At larger genome coverage (2%), F-Seq2 PE without filter mode, and SE mode showed superior performance (both have equal highest F-0.5 score = 0.62 at different coverages) versus all other modes. This observation suggests that the additional data improved precision for medium ranked peaks in F-Seq2 in its non-filter-based mode, which takes advantage of the greater genomic information available for more robust and accurate background estimations at the cost of precision at low genome coverage.

**Figure 2.**
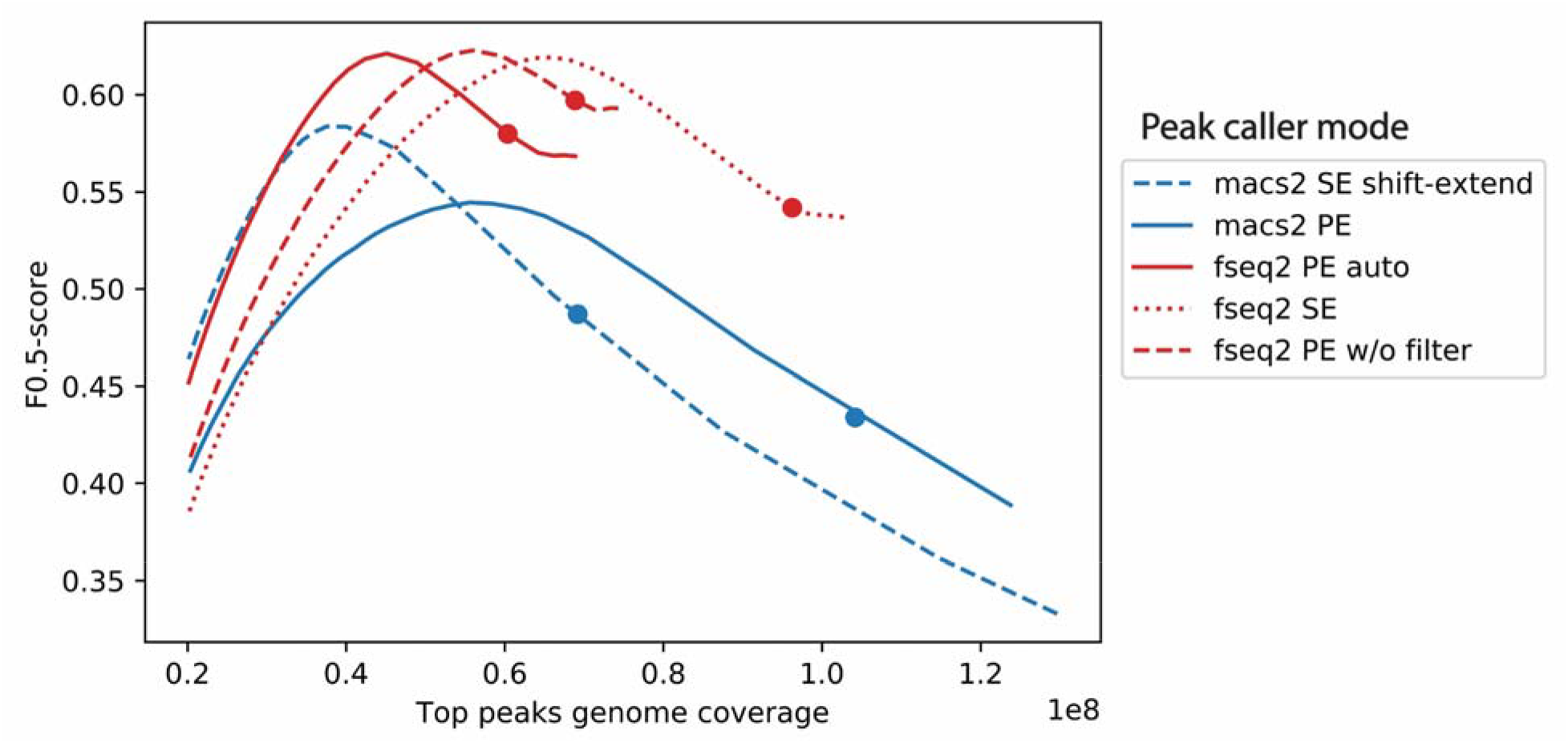
Comparison of F-Seq2 and MACS2 on the ATAC-seq paired-end data in GM12878. The F0.5-score operating characteristic curve where F0.5-score is plotted as a function of the genome coverage in base pairs by the top ranked peaks. F0.5-score put more emphasis on precision than recall due to the incompleteness of our “ground truth”. MACS2 was run with two modes: SE mode and PE mode. SE mode was run with shift (−75 bp) and extend (150 bp), which is used by the ENCODE ATAC-seq data analysis pipeline (12). F-Seq2 was run with three modes: PE auto mode, PE without filter mode, and SE mode. PE auto mode used the F-Seq2 auto filter which is designed based on nucleosome-related fragment length information (See Material and Methods for design details). Dots on curves indicate the genome coverage of significant peaks by the default threshold of each peak caller.

F-Seq2 was benchmarked on 3 real ChIP-seq datasets to confirm that the observed high performance under the simulated situations can be recapitulated using real data. F-Seq2 had the largest fraction of top *n* peaks (up to 1000 peaks) within 100 bp of a CTCF motif (Figure 3). GEM was the second largest with a slightly better performance than MACS2. The empirical distribution of the distance of called peaks to CTCF motifs showed that GEM stood out: 80% of its 1000 most significant peaks were within 4 bp of a CTCF motif, compared to F-Seq2 and MACS2 that were within 30 bp. This performance gain is due to GEM’s additional motif discovery utility allowing it to call peaks near discovered motifs. Similar trends were observed in the MAFK ChIP-seq data benchmarking results, the exception being SPP which had the most variable number of peaks called by a default threshold between the two TFs (Supplementary Figure 1). However, all peak callers had much lower and barely distinguishable performance on STAT1 (Supplementary Figure 2). (24) showed that 76 out of 220 chromatin factor ChIP-seq peaks lacked relevant sequence motifs, and STAT1 peaks were low in motif occupancy (below 50%), suggesting that evaluating peak callers using motifs may not reflect actual performance. The likely problematic evaluation with motifs renders it necessary to assess performance with the ground truth of simulated data, which is more accurate and precise.

**Figure 3.**
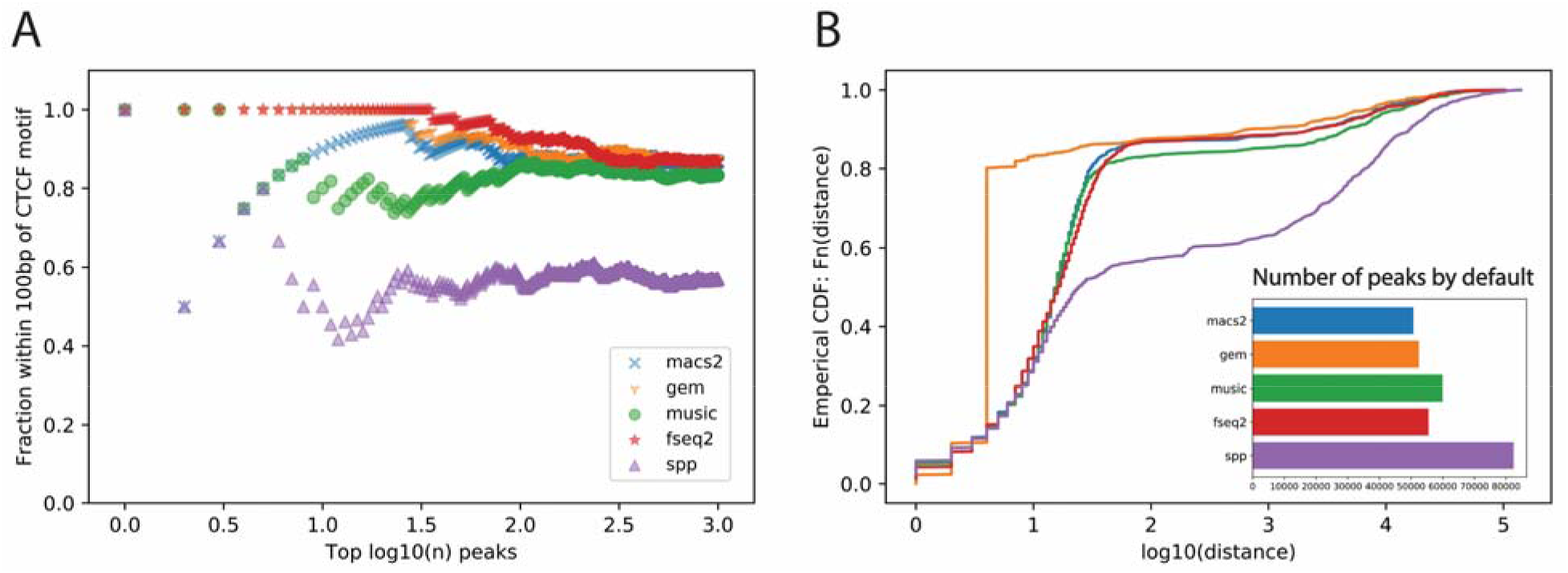
Comparison of peak callers on the CTCF ChIP-seq in ascending aorta female adult (51 years). (A) The fraction of top *n* peaks within 100 bp of a CTCF motif. (B) The empirical distribution of the shortest distance of the called peaks to a CTCF motif. The subplot shows the number of significant peaks called by each method using the default threshold.

## DISCUSSION

The highly-balanced performance of F-Seq2 between precision and recall across different assays is noteworthy. Kernel density estimation, which is a non-parametric method to model the read probability distribution, has an advantage over explicit modeling methods. Confounding experimental and biological factors, such as antibody specificity, DNA susceptibility to enzymes, and sequencing reads mappability, are difficult to form explicit assumptions (25), especially across different assays. The advantage of KDE has been demonstrated by the original peak caller F-Seq, which is the top-performing peak caller on DNase-seq datasets (1), and frequently used for FAIRE-seq data peak calling (3). We designed a new statistical framework and introduced new features to F-Seq to further improve the performance in its second version. Adding support of user-input control data allows for F-Seq2 to more accurately model background probability distribution together with the treatment reads distribution. With the help of a dynamic parameter, two local distributions around candidate summits can be summarized into significance values, leading to statistically robust peak ranks and peak calls. The joint effect of KDE and the dynamic parameter demonstrated superior performance in our benchmarking results, especially without control data. This suggests control information can be extracted from treatment data, given control and treatment data are well correlated. The support of control data allows for a more biologically meaningful signal to be reconstructed by weighting the treatment with control data, which leads to a better sanity-check when comparing and combining signals from different datasets (11).

Whether control data is a dispensable dataset for ChIP-seq peak calling requires further investigation. Recent papers (14, 26) that predict the linear weights for control datasets from treatment datasets provide evidence that control information can be extracted from treatment data. In our simulation results, a comparable performance was observed when using or omitting control data. F-Seq2 runs using experiments with real ChIP-seq data showed only a slightly decrease in performance without control data (data not shown). We suspect that the high correlations between control and treatment data explain the observation that control data is not required in a simulation setting. However, conclusions cannot be made based on the small performance difference on the real ChIP-seq datasets due to evaluation biases with motifs. We are unable to determine if a large observable discrepancy (low correlation) between control and treatment data is due to the low quality of either of the data, or to the indispensable information in control dataset.

F-Seq2 is compatible and suitable for IDR analysis which we recommend as a more reliable approach to determine a significance threshold when working with replicates. The IDR algorithm requires peak callers to run at a relaxed threshold to include both signal and noise peaks within the output to detect the consistency transition point between the two groups (15). During benchmarking, the MACS2 peak width detection was observed to be tied to peak detection. Specifically, when lowering the q-value threshold, by default MACS2 called not only more peaks, but larger width peaks, and may cause irreproducibility as a side-effect (i.e. changing the significance scores and ranks of called peaks). We developed F-Seq2 with summit-focused statistical testing and used separate parameters for peak width detection and summit detection. F-Seq2 reliably reproduces the exact summits and peaks, even when lowering the p-value or q-value threshold, and an individual significance score for each summit is calculated. Having separate scores for each summit and less rank ties by p-value interpolation are essential for the IDR to precisely identify the transition point, representing the desired threshold. We have built a peak calling pipeline for a pair of replicates with F-Seq2 followed by an integrated IDR analysis with our recommended settings, and is directly accessible through the command line interface.

F-Seq2 further pushes the potential in the mature field of peak calling. The accuracy of peak calling is essential for downstream analysis, such as differential analysis to discover new biological insights and mechanisms with HTS data.

## AVAILABILITY

### Data accessibility and peak caller parameter settings

Simulated data was reproduced from (16). The adapted scripts to simulate ChIP-seq data, and the scripts to run all peak callers are available at https://github.com/nsamzhao/F-Seq2-Paper-Supplementary. The accession numbers of all ENCODE data, and the IDs of all JASPAR motifs used in this study are also available at this website.

### Software availability

The F-Seq2 software and documentation are available at https://github.com/Boyle-Lab/F-Seq2. F-Seq2 can be installed through the Python Package Index (PyPI) and the Conda package manager.

## SUPPLEMENTARY DATA

Supplementary Data are available at NAR online.

## ACKNOWLEDGEMENT

We would like to thank members of the Boyle lab for critical reading and suggestions on the manuscript.

