## Supplemental Information for "F-Seq2: improving the feature density based peak caller with dynamic statistics"

F-Seq2 Supplementary Materials

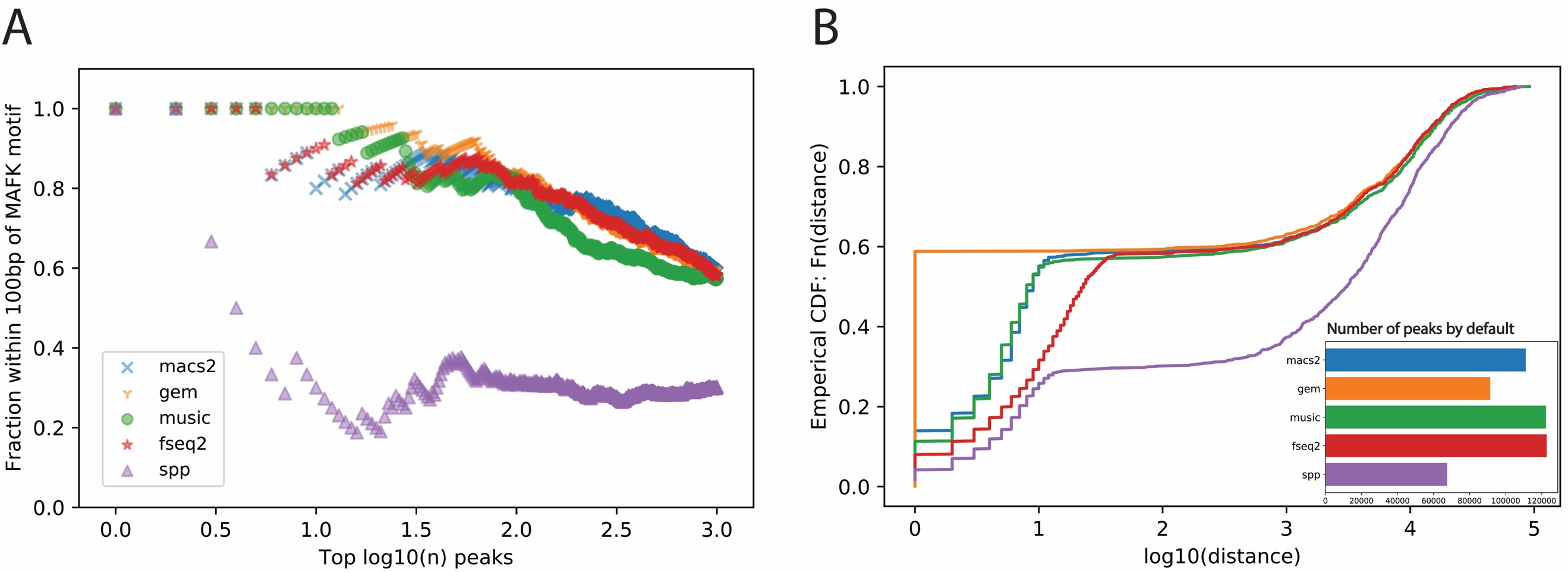

**Supplementary Figure S1. Comparison of peak callers on the MAFK ChIP-seq in HepG2.** (A) The fraction of top $n$ peaks within 100 bp of a MAFK motif. (B) The empirical distribution of the shortest distance of the called peaks to a MAFK motif. The subplot shows the number of significant peaks called by each method using the default threshold.

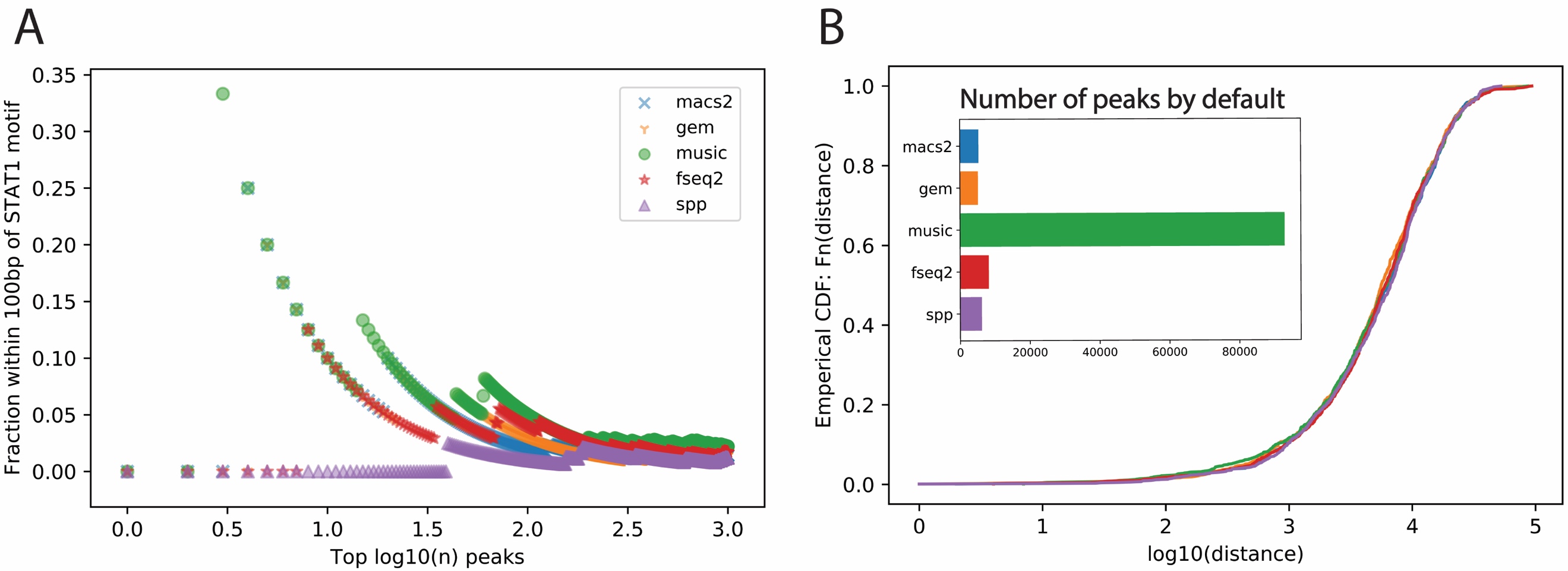

**Supplementary Figure S2. Comparison of peak callers on the STAT1 ChIP-seq in GM12878.** (A) The fraction of top $n$ peaks within 100 bp of a STAT1 motif. (B) The empirical distribution of the shortest distance of the called peaks to a STAT1 motif. The subplot shows the number of significant peaks called by each method using the default threshold.
